# Effects of breeding history and crop management on the root architecture of wheat

**DOI:** 10.1101/2020.02.25.964338

**Authors:** N. Fradgley, G. Evans, J.M. Biernaskie, J. Cockram, E.C. Marr, A. G. Oliver, E. Ober, H. Jones

## Abstract

**Aims:** Selection for optimal root system architecture (RSA) is important to ensure genetic gains in the sustainable production of wheat (*Triticum aestivum* L.). Here we examine the idea that past wheat breeding has led to changes in RSA and that future breeding efforts can focus directly on root traits to improve adaptation to a target environment.

**Methods:** We conducted three field trials using diverse wheat varieties, including modern and historic UK varieties and non-UK landraces, tested under contrasting tillage regimes (non-inversion tillage versus conventional ploughing) or different seeding rates (standard rate versus high rate). We used field excavation, washing and measurement of root crowns (‘shovelomics’) to characterise RSA traits, including: numbers of seminal, crown and nodal roots per plant, and crown root growth angle.

**Results:** We found large differences among genotypes for all root traits. Modern varieties generally had fewer roots per plant than historic varieties. There were fewer crown roots and root angles were wider, on average, under shallow non-inversion tillage compared with conventional ploughing. Crown root numbers per plant also tended to be smaller at a high seeding rate compared with the standard rate. There were significant genotype-by-year, genotype-by-tillage and genotype-by-seeding-rate interactions for many root traits.

**Conclusions:** Smaller root systems is likely to be a result of past selection and may have facilitated historical yield increases by reducing below-ground competition within the crop. The effects of crop management practices on RSA depend on genotype, suggesting that future breeding could select for improved RSA traits in resource-efficient farming systems.

## 1. Introduction

Increasing global human population growth, combined with challenges due to climate change and resource depletion, means that agriculture must become more productive and efficient while also contributing fewer greenhouse gas emissions (Conijn et al., 2018; Smith et al., 2007). Therefore, crop resource-use efficiency and adaptation to resource-efficient farming systems are key targets for crop genetic improvement. Wheat (*Triticum aestivum* L.) is a particularly important source of human and animal nutrition across the world (Shiferaw et al., 2013), so genetic improvements in the sustainable production of wheat would contribute greatly to the emerging challenges in global food security.

An underappreciated route to more productive and efficient wheat crops is via genetic improvements in root system architecture (RSA). Evidence suggests that RSA is integral to crop nutrient uptake, water acquisition and grain yield (Smith and De Smet, 2012) and that changes in RSA are linked to historical improvements in wheat productivity (Zhu et al. 2019a). It has been suggested that targeting RSA for crop improvement could lead to a second Green Revolution, where increased resource capture could further enhance yields and reduce the need for fertiliser (Lynch, 2007). However, plant breeders have largely neglected direct selection for wheat root traits. This is in part due to the relative inaccessibility of roots, their phenotypic plasticity, and the absence of high-throughput screening methods (Manschadi et al., 2006). Current root phenotyping methods have mostly focused on root traits in young plants under controlled environments (Atkinson et al., 2015; Kuijken et al., 2015; Richard et al., 2015; Watt et al., 2013). However, these techniques do not reflect real soil conditions in the field, and inconsistent results are often found between methods (Wojciechowski et al., 2009). On the other hand, current RSA phenotyping methods in field conditions are slow, laborious and prone to excessive variation (Gregory et al., 2009).

Improved RSA phenotyping would be particularly useful in field conditions that reflect resource-efficient farming systems. In developing countries, crop productivity is often limited by soil erosion and by access to inputs such as fertilisers, whereas in high-input systems, inefficient use of inputs by the crop can result in unused nutrients (e.g., nitrogen and phosphorus) causing environmental damage (Ascott et al., 2017; Cordell et al., 2009; FAO, 2016). Low-input agriculture may benefit from the principles of conservation agriculture which include tillage practices that minimise soil disturbance and provide several environmental benefits (Hobbs, 2007; Mangalassery et al., 2014; Petersen et al., 2008), promotion of soil microbial activity (Kabir, 2005; Papp et al., 2018), and improved soil structure which limits soil erosion (Zhang et al., 2007). Relatively high-input agriculture, on the other hand, could benefit from high-density cropping systems, where crops with higher plant density may collectively make better use of the available nutrients (Donald, 1968; Marin and Weiner 2014). Plant breeding and evaluation of different crop varieties, however, are rarely conducted under the conditions of conservation agriculture or high-density cropping.

To address these issues, we use a semi-high-throughput, field-based method of phenotyping wheat RSA traits in the context resource-efficient farming systems. Our approach involves field excavation, washing and measurement of root crowns (‘shovelomics’; Trachsel et al., 2011; Burridge et al., 2016; Colombi et al., 2015; York et al., 2018), and uses modern and historic UK wheat varieties and non-UK landraces, tested under contrasting tillage regimes (non-inversion tillage versus conventional ploughing) or different seeding rates (standard rate versus high rate). We investigate the idea that past wheat breeding has led to consistent changes in RSA and that future breeding efforts can focus directly on root traits to improve adaptation to a target environment. Specifically, our aims are to examine: (1) how wheat RSA traits vary with their variety’s year of release; and (2) how wheat RSA traits respond to changes in tillage regime or seeding rate and whether genotypes vary in these responses.

## 2. Materials and Methods

### 2.1. Germplasm

The genotypes from two panels of wheat cultivars were chosen to represent a wide range of diversity, including modern, historic and landrace accessions.

The WHEALBI panel consisted of 20 UK and non-UK modern and historic wheat genotypes. Ten lines were a subset of the larger WHEALBI panel (Pont et al. 2019), and ten additional lines were chosen by collaborators at the Organic Research Centre (Supp. table 1). Seed for UK historic cultivars and non-UK landrace accessions was sourced from the John Innes Centre Germplasm Resource Unit in the UK (GRU http://www.jic.ac.uk/germplasm/). Five non-UK landrace accessions were chosen from the full Watkins collection, which consists of 826 landrace accessions originating from a wide range of non-UK backgrounds (Wingen et al., 2014). Hungarian lines were supplied by ATK (Hungary), and Tiepolo was supplied by SIS (Italy). Seed stocks were multiplied in 1 m^2^ nursery plots at NIAB, Cambridge in 2014/15. Seed from currently grown modern varieties was sourced from seed merchants.

The 16 founders of a multi-founder advanced generation inter-cross (MAGIC) population (‘NIAB Diverse MAGIC’) were chosen to capture the greatest genetic diversity based on genetic markers from the set of 94 UK and northern European wheat varieties described in White et al. (2008) (Supp. Table 2). Seed was used from stock maintained at NIAB but originally sourced from the John Innes Centre Germplasm Resource Unit.

### 2.2. Field trial sites

Autumn-sown field trials were carried out at two sites. The WHEALBI panel of 20 accessions was grown over two trial years (Autumn 2015 to Summer 2016 and Autumn 2016 to summer 2017) at Reading University research farm, Sonning, Berkshire, UK (Lat: 51.481470, Long: -0.89969873). The 16 NIAB Diverse MAGIC founders were trialled in one trial year (Autumn 2017 to Summer 2018) at Duxford, Cambridgeshire, UK (Lat: 52.099091, Long: 0.13352841). The soil at the Sonning site was classified as a Luvisol and described as a loam over gravel. The soil chemistry was measured at drilling and is summarised in Supp. table 3. In each year, the trial was located on a different field section at the same site. The total precipitation was 535 and 575 mm for the growing seasons in year 1 and 2, respectively. The soil at the Duxford trial site was a freely draining lime-rich loam and total precipitation for the season was 359 mm.

### 2.3. Trial design and management

#### 2.3.1. Sonning trial site

The trial site was managed under organic farming practices and the trials were conducted in the first cereal position in the rotation following a two-year grass ley (comprising cocksfoot, red clover, white clover and black medic). Trials were conducted using the 20 winter wheat genotypes from the WHEALBI panel in a split plot design, with tillage treatments as main plots, and cultivar as sub-plots with four replications. Cultivars were randomised within each block. Transition areas between tillage treatments were sown with discard crop plots to minimise edge effects. Tillage treatments were conventional plough tillage (CT) to a depth of 250 mm and shallow non-inversion tillage (SNI), performed using a shallow rotovator (50-75 mm depth). In both treatments, seedbeds were prepared with a power harrow set to 125 mm depth. In the CT treatment, the previous ley was mown before ploughing to a depth of 250 mm, whilst in SNI, the ley was terminated using a rotovator at a depth of 50-75 mm. A power harrow was used to create a seedbed in both cultivation systems before sowing seeds using a plot direct drill with front discs. The plots were sown on 12/10/2015 and 02/11/2016 in years 1 and 2, respectively. Trial plots consisted of 14 rows 15 cm apart so that plot dimensions were 2.1 m wide and 7.5 m long. Seed rates were adjusted to achieve a target plant population of 500 plants m^-2^ taking into account seed weight and germination rate. Plots were rolled to consolidate the seedbed after drilling. Mechanical weeding was carried out using a spring tine harrow in year 2 as required but this could not be used in year 1 due to high rainfall. Seeds were treated with 10 g/kg of Tillecur® (yellow mustard powder; Biofa AG, Germany) plant strengthening seed treatment to control common bunt and other seed-borne diseases.

#### 2.3.2. Duxford trial site

The Duxford site was managed conventionally. Fertiliser inputs included 110 kg ha^-1^ of nitrogen in the form of prilled ammonium nitrate over three timings in February, April and May. This was at half the field recommended rate to manage lodging risk in tall varieties. Herbicides were used to control grass and broad-leafed weeds in November and in May. Fungicides were used to control foliar diseases applied at three timings from April to June and plant growth regulators were applied in April and May to reduce lodging risk. Insecticide was applied in June to control orange wheat blossom midge. Seeds were treated with systemic fungicide to control seed-borne diseases. Four plot replicates of each cultivar from the NIAB Diverse MAGIC founder panel were sown at two sowing rates (300 [a standard rate of local practice] and 600 plants m^-2^), after adjusting for mean seed weight. Plots were randomised within a larger trial of 2,380 plots of the full MAGIC population. Plots were sown over two days on the 13/10/2017 and 14/10/2017 and consisted of 12 rows 14 cm apart so that plot dimensions were 1.54 m wide and 6 m long. The field was ploughed before cultivations to create a seedbed before sowing.

### 2.3. Crop assessments

Root samples were taken on 14/07/2016 and 20/07/2017 in year 1 and year 2, respectively, at Sonning, and on 01/08/2018 at Duxford when the crop was at approximately growth stage GS80 (Zadoks et al., 1974). At both sites, two samples, including the base of the crop plant, roots and surrounding soil, were taken per plot using a 20 cm wide and 30 cm deep shovel, bagged and stored before analysis. This method ensured that the position and integrity of the roots within this volume were not affected while in storage.

Root samples were processed by soaking each sample in water with detergent for approximately five minutes before manually washing the soil from the crop roots and plant base. A randomly chosen single plant was taken per sample for scoring root traits. Samples from trials at the Sonning site in 2016 and 2017 were imaged and later scored from a digital image whereas samples from Duxford in 2018 were manually scored *in situ* directly after washing. Images were taken against a dark background using a Canon EOS 1000 digital camera with F-stop set to f/25, exposure time at 1/4 second and ISO at 200. Two images were taken per sample changing the orientation by 90° in the second image. Each sample was then divided into their constituent tillers (including adjoining roots), and each tiller individual was imaged at two 90° orientations. Digital images were subsequently used to visually score root traits. ImageJ2 image analysis software (Rueden et al., 2017) was used to manipulate images and improve contrast for scoring. The RSA traits scored were root angle (RA), crown root number (CRN), nodal root number (NRN) and seminal root number (SRN), as detailed further in Table 2 and illustrated in Figure 1. It was only possible to measure SRN on 88% of the samples from images in the Sonning dataset due to the coleoptile and seed growing point often being obscured in the image. Harvest grain yield at the Duxford site was determined using a small plot combine and yields were adjusted to 15 % moisture content.

**Table 2.** Description of wheat root traits scored from imaged shovelomics samples.

| Trait | Abbreviation | Description |
| --- | --- | --- |
| Root angle | RA | The angle between two lines originating at the base of the plant at ground level which fits the angle of the majority of the crown roots in a 2D image of the whole plant using the angle tool function within ImageJ (Rueden et al., 2017) analysis software (Figure 1). |
| Crown root number | CRN | Number of roots originating from the base of the plant at ground level. |
| Nodal root number | NRN | The number of roots originating from the first node. |
| Seminal root number | SRN | The number of roots originating from the germinated seed below the coleoptiles. |

**Figure 1.**
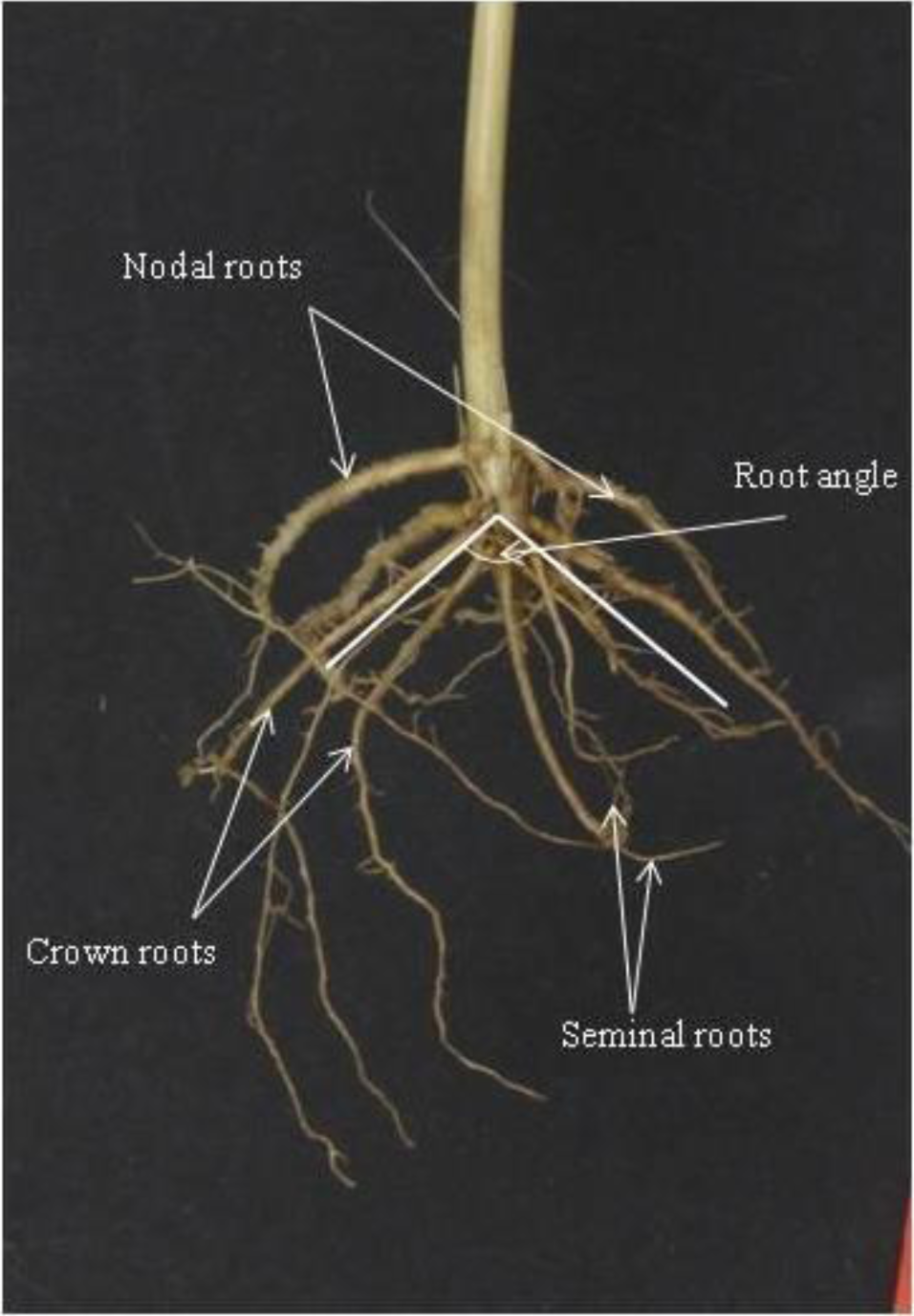
Example image of a wheat root sample obtained using the shovelomics methodology with annotations of root phenotypes scored.

### 2.4. Statistical analysis

All data analyses were carried out using Genstat (18^th^ edition) statistical analysis software (Payne et al., 2009). Plot data used in these analyses are available in the Supplementary Table 4. Data from each cultivar panel at Duxford and Sonning were analysed separately. Both trial years at Sonning were combined for the analyses at this site. Data for RA and SRN were analysed using Linear Mixed Effects Models (LMMs) whilst count data with non-normally distributed residuals for CRN and NRN were analysed using Generalised Linear Mixed Effects Models (GLMMs), including Poisson error structure and logarithmic link function with dispersion fixed to one. For both trial years from Sonning, year, tillage and genotype were considered interacting fixed effect terms in that order, whilst blocks nested within year and blocks within tillage within year were considered as random effects in both LMMs and GLMMs. For data from Duxford, genotype and sowing rate were included as interacting fixed effects and main experimental block was included as a single random effect. For fixed effects, model simplification from the maximal model was performed based on the Wald test for GLMMs and F statistic for LMMs where non-significant terms (p>0.05) were removed. Random effect terms were removed when negative variance components were found. Adjusted genotypic predicted mean values were calculated for each trait as generalised means across fixed effects. Then, where significant interacting fixed effect terms were found, separate models were run for each interacting term level, and deconstructed adjusted genotypic mean values were also calculated separately for interacting factor levels when significant effects of genotype were found (p<0.05). Correlations among generalised varietal adjusted mean phenotypic values, as well as genotype by year of release were determined using the Pearson correlation coefficient.

## 3. Results

### 3.1. Genotypic differences and trends in root architecture over time

Wheat RSA traits were phenotyped using the shovelomics method using two diverse sets of wheat varieties in multiple environments. Generalised analysis of these data across both years and fixed effects revealed statistically significant genotypic differences for all studied root phenotypes examined in the sets of varieties at both the Sonning (Table 3) and Duxford sites (Table 4). Differences amongst genotypes were significant for RA and highly significant for CRN and NRN in both datasets. A highly significant genotype effect was found for SRN among the 16 NIAB Diverse MAGIC founders grown at Duxford and among the 20 WHEALBI accessions grown at Sonning. The consistency of these traits was also be compared between the two datasets where three varieties (‘Steadfast’, ‘Robigus’ and ‘Soissons’) were in common. The ranking of these three varieties was consistent for CRN and NRN, with ‘Steadfast’ having the greatest CRN and NRN. However, rankings for RA and SRN between these three varieties were not consistent, indicating stronger genotype-by-environment interactions for these traits.

**Table 3.**
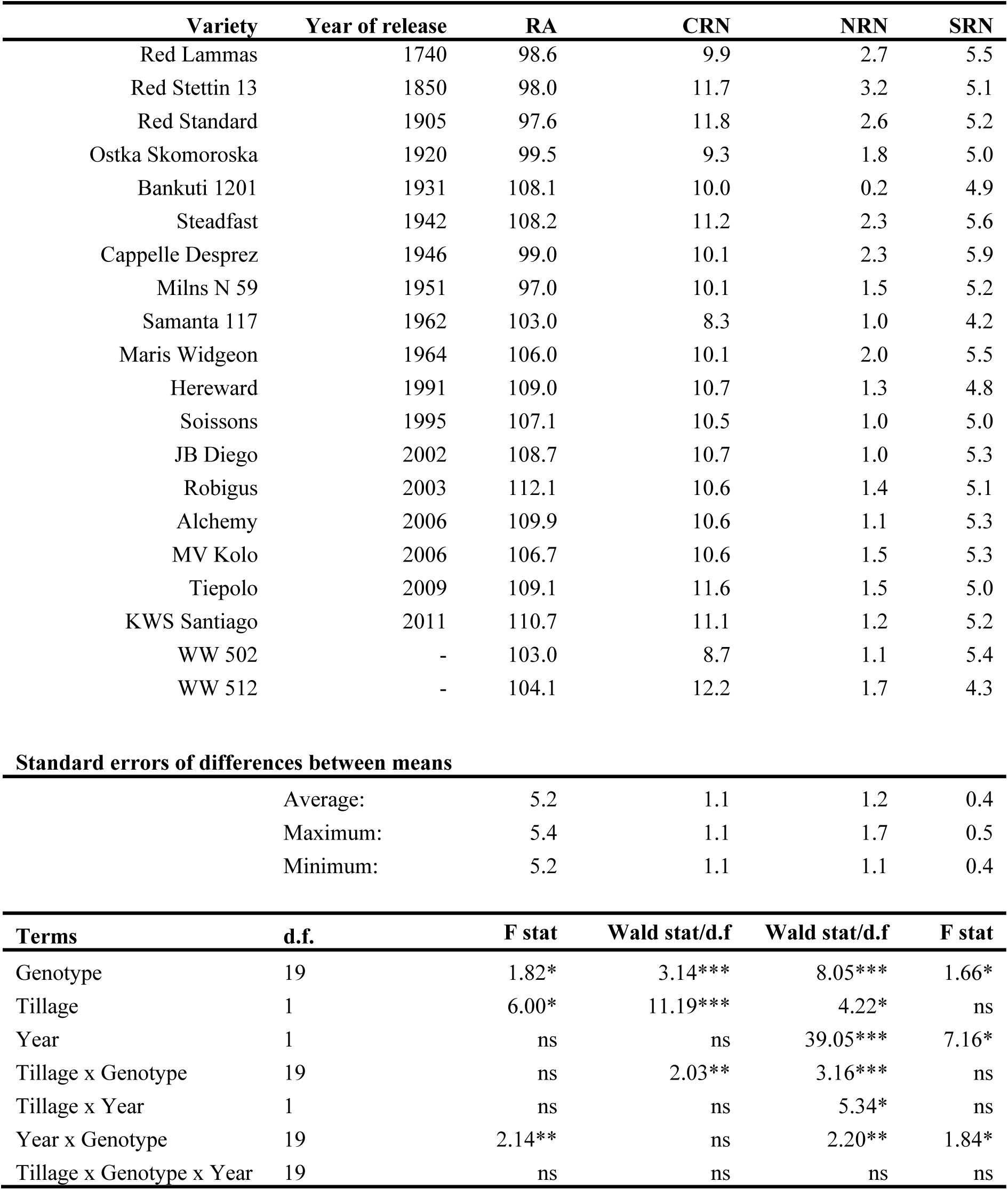
Generalised predicted mean values among 20 wheat varieties for all root traits generalised across two tillage levels and over two trial years at the Sonning trial site. RA = root angle, CRN = crown root number, NRN = nodal root number, SRN = seminal root number. Asterisks indicate significance level: *** = p<0.001, ** = p<0.01 *, = p<0.05, ns = not significant. d.f. = degrees of freedom.

| Variety | Year of release | RA | CRN | NRN | SRN |
| --- | --- | --- | --- | --- | --- |
| Red Lammas | 1740 | 98.6 | 9.9 | 2.7 | 5.5 |
| Red Stettin 13 | 1850 | 98.0 | 11.7 | 3.2 | 5.1 |
| Red Standard | 1905 | 97.6 | 11.8 | 2.6 | 5.2 |
| Ostka Skomoroska | 1920 | 99.5 | 9.3 | 1.8 | 5.0 |
| Bankuti 1201 | 1931 | 108.1 | 10.0 | 0.2 | 4.9 |
| Steadfast | 1942 | 108.2 | 11.2 | 2.3 | 5.6 |
| Cappelle Desprez | 1946 | 99.0 | 10.1 | 2.3 | 5.9 |
| Milns N 59 | 1951 | 97.0 | 10.1 | 1.5 | 5.2 |
| Samanta 117 | 1962 | 103.0 | 8.3 | 1.0 | 4.2 |
| Maris Widgeon | 1964 | 106.0 | 10.1 | 2.0 | 5.5 |
| Hereward | 1991 | 109.0 | 10.7 | 1.3 | 4.8 |
| Soissons | 1995 | 107.1 | 10.5 | 1.0 | 5.0 |
| JB Diego | 2002 | 108.7 | 10.7 | 1.0 | 5.3 |
| Robigus | 2003 | 112.1 | 10.6 | 1.4 | 5.1 |
| Alchemy | 2006 | 109.9 | 10.6 | 1.1 | 5.3 |
| MV Kolo | 2006 | 106.7 | 10.6 | 1.5 | 5.3 |
| Tiepolo | 2009 | 109.1 | 11.6 | 1.5 | 5.0 |
| KWS Santiago | 2011 | 110.7 | 11.1 | 1.2 | 5.2 |
| WW 502 | - | 103.0 | 8.7 | 1.1 | 5.4 |
| WW 512 | - | 104.1 | 12.2 | 1.7 | 4.3 |

**Standard errors of differences between means**
|  |  |  |  |  |
| --- | --- | --- | --- | --- |
| Average: | 5.2 | 1.1 | 1.2 | 0.4 |
| Maximum: | 5.4 | 1.1 | 1.7 | 0.5 |
| Minimum: | 5.2 | 1.1 | 1.1 | 0.4 |

| Terms | d.f. | F stat | Wald stat/d.f | Wald stat/d.f | F stat |
| --- | --- | --- | --- | --- | --- |
| Genotype | 19 | 1.82* | 3.14*** | 8.05*** | 1.66* |
| Tillage | 1 | 6.00* | 11.19*** | 4.22* | ns |
| Year | 1 | ns | ns | 39.05*** | 7.16* |
| Tillage x Genotype | 19 | ns | 2.03** | 3.16*** | ns |
| Tillage x Year | 1 | ns | ns | 5.34* | ns |
| Year x Genotype | 19 | 2.14** | ns | 2.20** | 1.84* |
| Tillage x Genotype x Year | 19 | ns | ns | ns | ns |

**Table 4.**
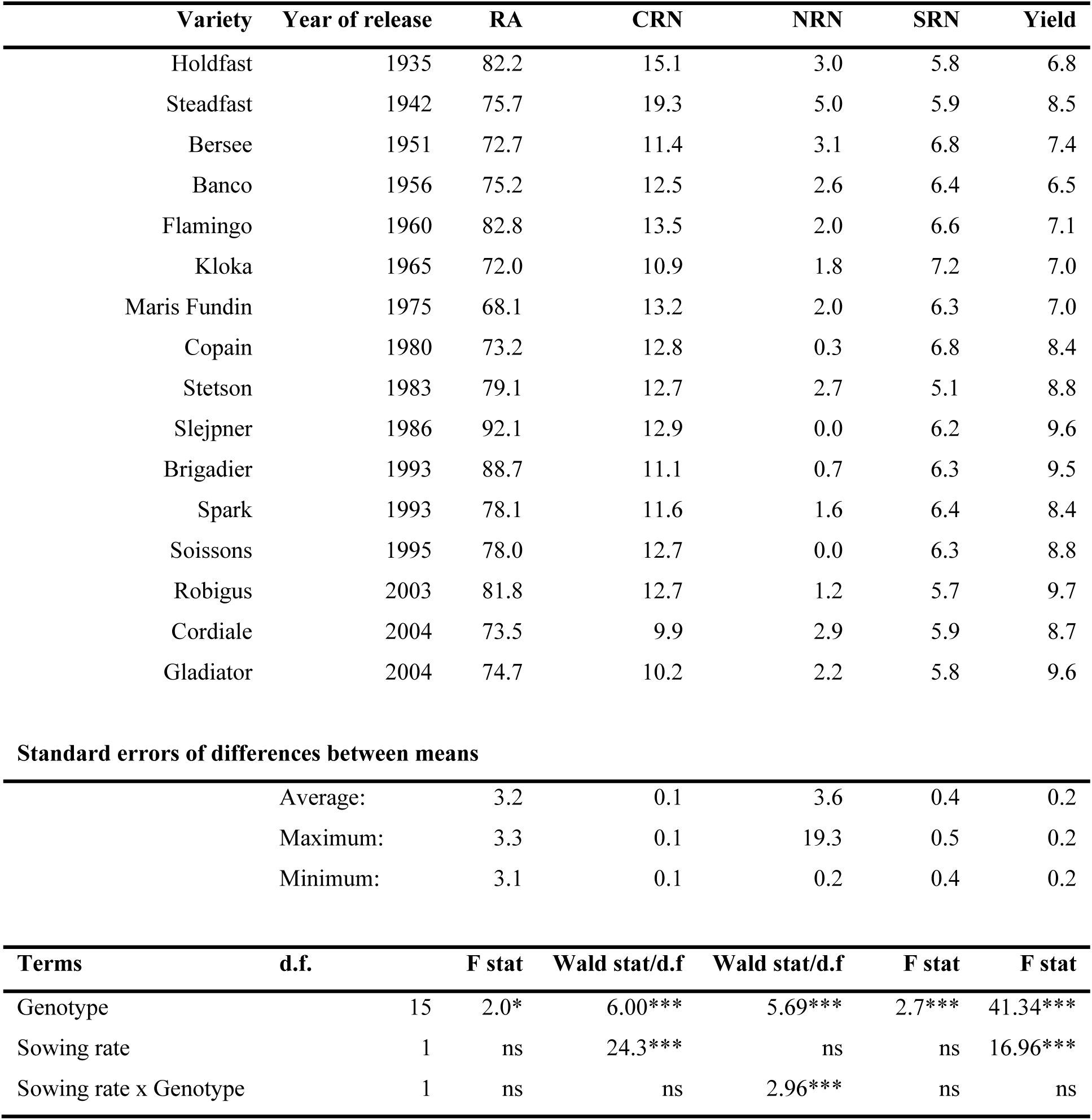
Generalised predicted mean values for all root traits across two seeding rate treatments and effects of experimental terms on root traits among 16 wheat varieties at the Duxford trial site in 2018. RA = root angle, CRN = crown root number, NRN = nodal root number and SRN = seminal root number. Asterisks indicate significance level: *** = p<0.001, ** = p<0.01 *, = p<0.05, ns = not significant. d.f. = degrees of freedom.

| Variety | Year of release | RA | CRN | NRN | SRN | Yield |
| --- | --- | --- | --- | --- | --- | --- |
| Holdfast | 1935 | 82.2 | 15.1 | 3.0 | 5.8 | 6.8 |
| Steadfast | 1942 | 75.7 | 19.3 | 5.0 | 5.9 | 8.5 |
| Bersee | 1951 | 72.7 | 11.4 | 3.1 | 6.8 | 7.4 |
| Banco | 1956 | 75.2 | 12.5 | 2.6 | 6.4 | 6.5 |
| Flamingo | 1960 | 82.8 | 13.5 | 2.0 | 6.6 | 7.1 |
| Kloka | 1965 | 72.0 | 10.9 | 1.8 | 7.2 | 7.0 |
| Maris Fundin | 1975 | 68.1 | 13.2 | 2.0 | 6.3 | 7.0 |
| Copain | 1980 | 73.2 | 12.8 | 0.3 | 6.8 | 8.4 |
| Stetson | 1983 | 79.1 | 12.7 | 2.7 | 5.1 | 8.8 |
| Slejpner | 1986 | 92.1 | 12.9 | 0.0 | 6.2 | 9.6 |
| Brigadier | 1993 | 88.7 | 11.1 | 0.7 | 6.3 | 9.5 |
| Spark | 1993 | 78.1 | 11.6 | 1.6 | 6.4 | 8.4 |
| Soissons | 1995 | 78.0 | 12.7 | 0.0 | 6.3 | 8.8 |
| Robigus | 2003 | 81.8 | 12.7 | 1.2 | 5.7 | 9.7 |
| Cordiale | 2004 | 73.5 | 9.9 | 2.9 | 5.9 | 8.7 |
| Gladiator | 2004 | 74.7 | 10.2 | 2.2 | 5.8 | 9.6 |

Standard errors of differences between means
|  |  |  |  |  |  |
| --- | --- | --- | --- | --- | --- |
| Average: | 3.2 | 0.1 | 3.6 | 0.4 | 0.2 |
| Maximum: | 3.3 | 0.1 | 19.3 | 0.5 | 0.2 |
| Minimum: | 3.1 | 0.1 | 0.2 | 0.4 | 0.2 |

| Terms | d.f. | F stat | Wald stat/d.f | Wald stat/d.f | F stat | F stat |
| --- | --- | --- | --- | --- | --- | --- |
| Genotype | 15 | 2.0* | 6.00*** | 5.69*** | 2.7*** | 41.34*** |
| Sowing rate | 1 | ns | 24.3*** | ns | ns | 16.96*** |
| Sowing rate x Genotype | 1 | ns | ns | 2.96*** | ns | ns |

Correlations among generalised predicted means across tillage or sowing rate treatments revealed clear trends in RSA over time (according to year varieties were released) as well as relationships among traits (Table 5). Modern varieties in both the datasets generally had fewer nodal roots than older cultivars (Figure 2). For example, the UK landrace variety ‘Red Stettin 13’ had more than twice as many nodal roots as any modern variety released after 1990 in the Sonning dataset. Only 31 % and 25 % of plants measured for the relatively modern varieties ‘Slejpner’ and ‘Soissons’ respectively, had any nodal roots at all in the Duxford dataset. The negative correlation between CRN and year of release was only significant at Duxford (Fig. 2b). A significant positive correlation between RA and year of release was observed only at Sonning, based on analysis across both years. This indicates that the spread of crown roots increased over time, with older varieties tending to have more narrow root systems. The relationship was perhaps more pronounced in the Sonning dataset because of the presence of old landraces with much narrower root RA than modern cultivars.

**Table 5.**
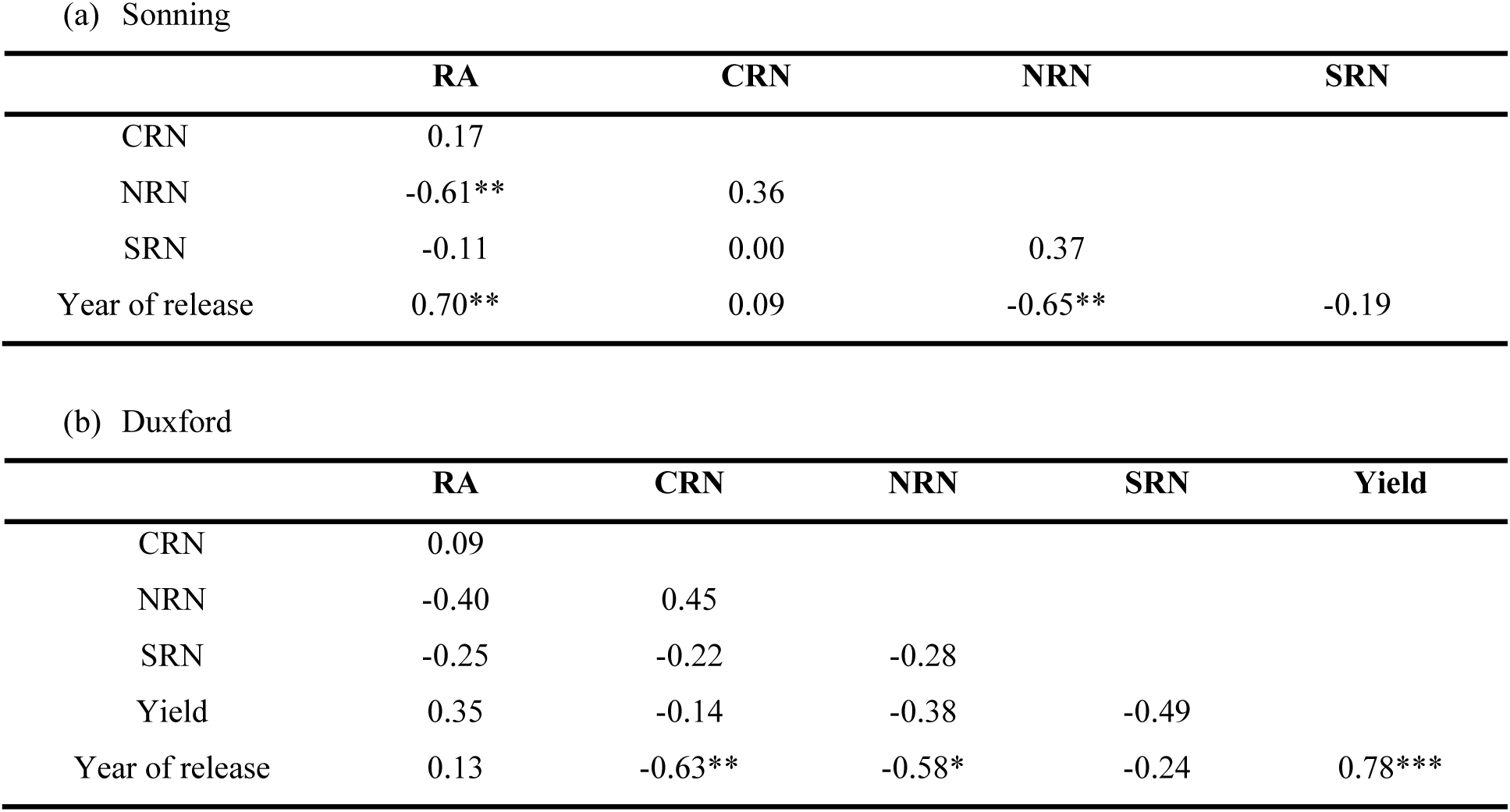
Correlation coefficients among predicted mean values for root traits for the set of 20 wheat varieties at the Sonning site (a) and the 16 wheat varieties at the Duxford Site (b). Root angle (RA), crown root number (CRN), nodal root number (NRN), seminal root number (SRN). Asterisks indicate significance level: *** = p<0.001, ** = p<0.01, * = p<0.05.

**Figure 2.**
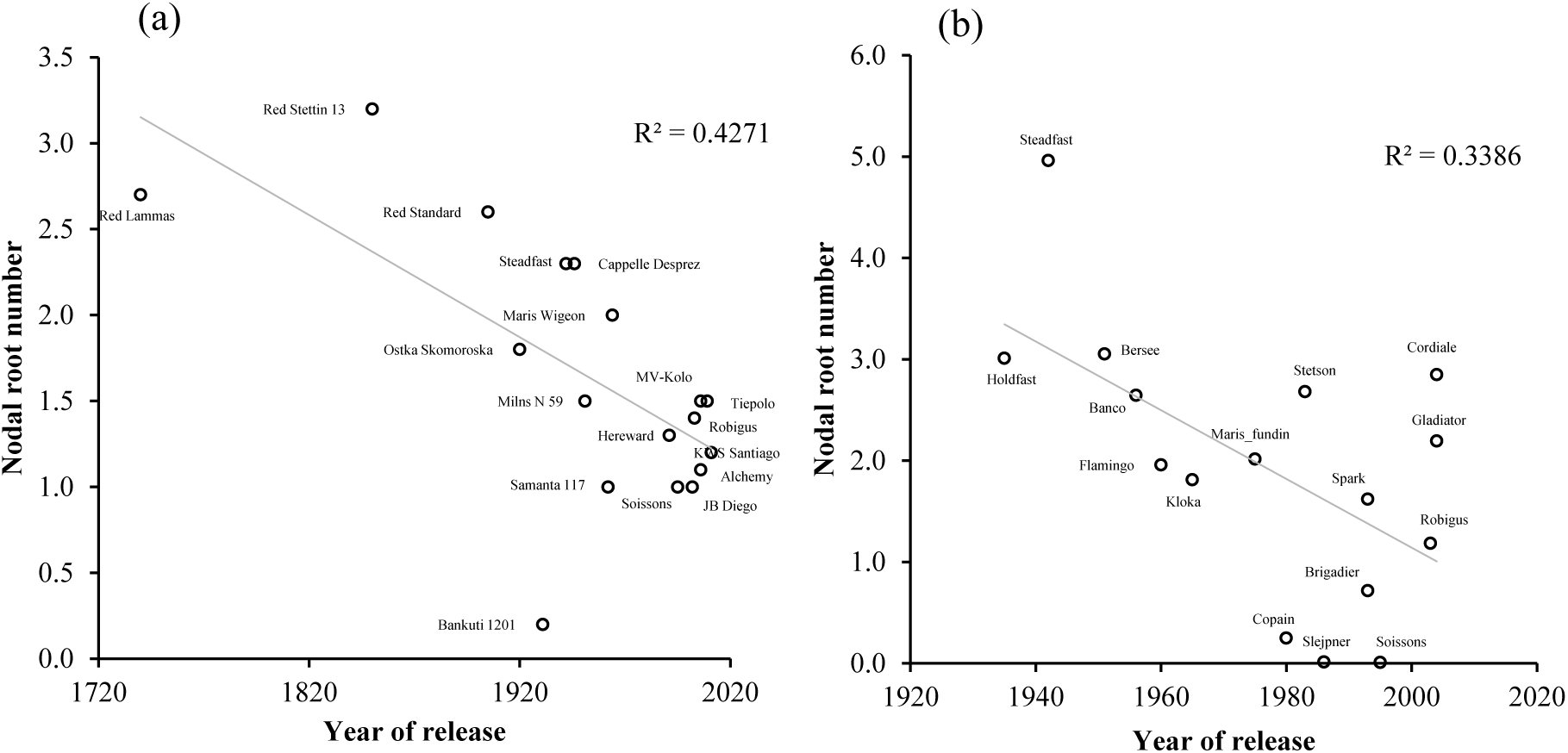
The relationship between year of varietal release and nodal root number for the set of 18 wheat varieties at the Sonning site with release date information (a) and the 16 wheat varieties at the Duxford Site (b).

### 3.2. Effects of tillage and genotype interactions

The dataset at Sonning allowed comparison of effects of contrasting inversion and shallow non-inversion tillage regimes as well as the variety response to these effects. All traits except SRN were affected by tillage (Table 3): there were generally more crown and nodal roots in conventional tillage (CT) and roots were at a wider angle than shallow non-inversion tillage (SNI).

RA in CT was on average 106.8° while in SNI the average root angle was 102.8°. As a significant genotype-by-year interaction was found for RA, further analysis was carried out separately for each year. This analysis showed the effect of genotype on RA was significant in the first year, where again RA was wider in CT than in SNI, but not significant in the second year (Table 6) when RA tended to be narrower in SNI (100.3°) than CT (105.6°) (Table 6). Although the genotype-by-year-by-tillage three-way interaction was not significant, the genotype-by-tillage interaction on RA was significant in year 1 (Table 6), where genotypic differences in RA were much more apparent in CT than under SNI (Table 6,7).

**Table 6.** Deconstruction of genotype-by-year interactions including effects of experimental terms on root angle (RA), nodal root number (NRN) and seminal root number (SRN) among 20 wheat varieties at two tillage levels at the Sonning site analysed separately for the two trial years. Effect values for size of each term include F-statistic for RA and SRN and Wald statistic/d.f. for NRN. Asterisks indicate significance level: *** = p<0.001, ** = p<0.01 *, = p<0.05 and ns” indicates non-significance. d.f. = degrees of freedom.

| Trait | Term | d.f. | Year 1 | Year 2 |
| --- | --- | --- | --- | --- |
| RA | Tillage | 1 | ns | 5.03* |
|  | Genotype | 19 | 3.36*** | ns |
|  | Tillage x Genotype | 19 | 2.05** | ns |
| NRN | Tillage | 1 | ns | 6.4* |
|  | Genotype | 19 | 3.46*** | 6.77*** |
|  | Tillage x Genotype | 19 | ns | 2.95*** |
| SRN | Tillage | 1 | ns | ns |
|  | Genotype | 19 | ns | 2.02* |
|  | Tillage x Genotype | 19 | ns | ns |

**Table 7.** Deconstruction of genotype-by-tillage interactions including effects of experimental terms on root angle (RA) in year 1, crown root number (CRN) in both years and nodal root number (NRN) in year 2 among 20 wheat varieties at the Sonning site analysed separately for two tillage levels. CT = conventional tillage and SNI = shallow non-inversion tillage. Effect values for size of each term include F-statistic for RA and Wald statistic/d.f. for CRN and NRN. Asterisks indicate significance level: *** = p<0.001, ** = p<0.01 *, = p<0.05. d.f. = degrees of freedom.

| Trait | Term | d.f. | CT | SNI |
| --- | --- | --- | --- | --- |
| RA in Year 1 | Genotype | 19 | 3.65*** | 1.85* |
| CRN in both years | Genotype | 19 | 1.93** | 3.23*** |
| NRN in Year 2 | Genotype | 19 | 4.48*** | 5.24*** |

The number of crown roots per plant was generally higher in CT (11.0) than SNI (10.0) across both years (Table 8). However, a small but significant genotype-by-tillage interaction was also found (Table 3). When each tillage system were analysed separately, the genotypic effect was greater in SNI than in CT (Table 7). In addition to the highly significant main effect of genotype on NRN, interactions of genotype-by-tillage, genotype-by-year and tillage-by-year were also found to be significant (Table 3). In the two cultivation systems tested, wheat grown under SNI (1.2) had fewer nodal roots per plant than under CT (1.7). There were more nodal roots per plant in the second year of trials (2.8) compared with the first year (1.1). When the two years were analysed separately, the effect of genotype was found to be highly significant in both years (Table 6). However, a significant genotype-by-tillage interaction was also found in year 2 (Table 6) where there were more crown roots in CT (3.5) than SNI (2.0). When CT and SNI were analysed separately in year 2, highly significant effects of genotype were found for NRN in both systems (Table 7).

**Table 8.**
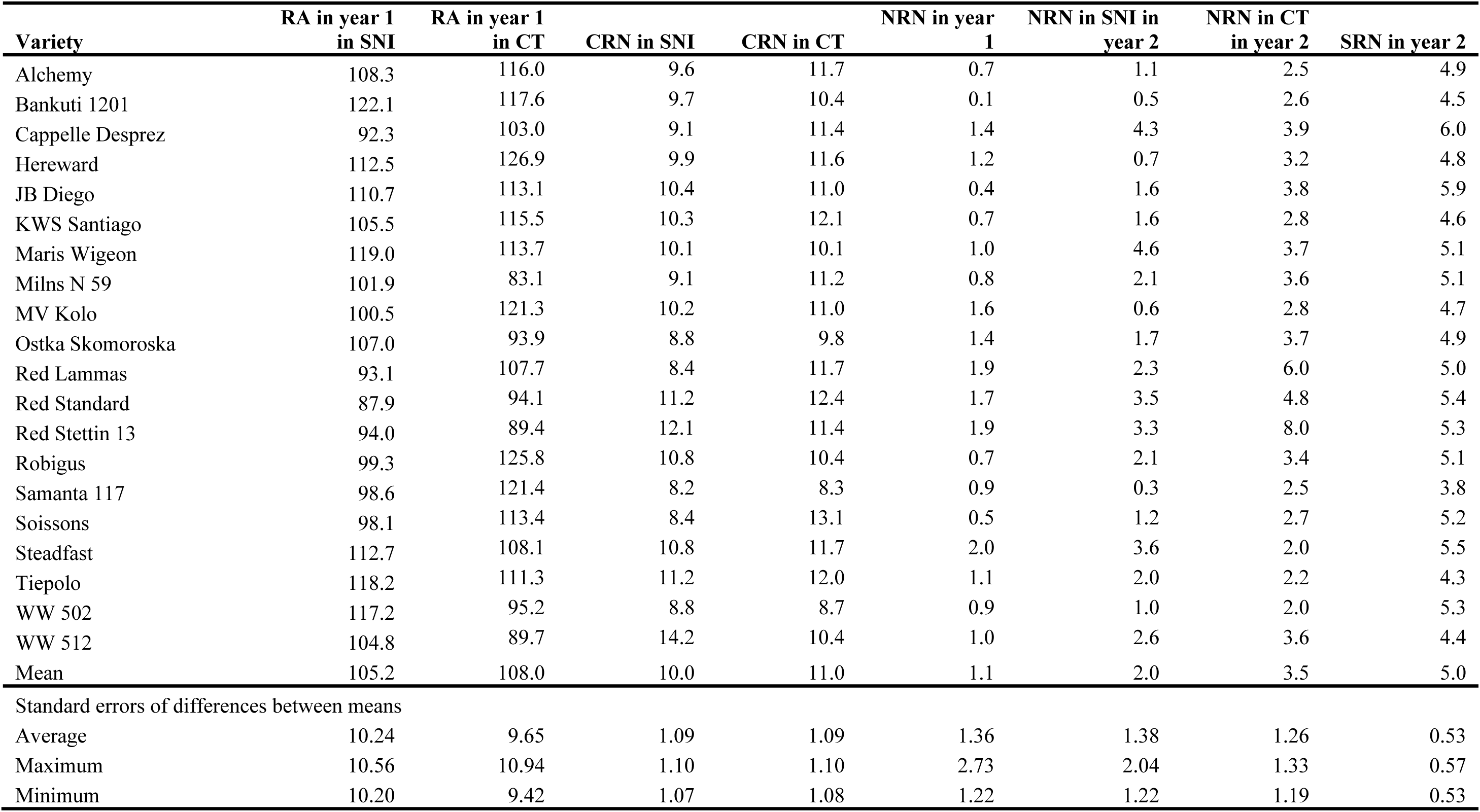
Predicted mean values after deconstruction of fixed effect interactions of root angle (RA), crown root number (CRN), nodal root number (NRN) and seminal root number (SRN) for 20 wheat varieties at the Sonning site. Means were calculated separately for different year or tillage levels where significant interactions with variety were found. Tillage levels include conventional tillage (CT) and shallow non-inversion tillage (SNI).

| Variety | RA in year 1<br>in SNI | RA in year 1<br>in CT | CRN in SNI | CRN in CT | NRN in year<br>1 | NRN in SNI in<br>year 2 | NRN in CT<br>in year 2 | SRN in year 2 |
| --- | --- | --- | --- | --- | --- | --- | --- | --- |
| Alchemy | 108.3 | 116.0 | 9.6 | 11.7 | 0.7 | 1.1 | 2.5 | 4.9 |
| Bankuti 1201 | 122.1 | 117.6 | 9.7 | 10.4 | 0.1 | 0.5 | 2.6 | 4.5 |
| Cappelle Desprez | 92.3 | 103.0 | 9.1 | 11.4 | 1.4 | 4.3 | 3.9 | 6.0 |
| Hereward | 112.5 | 126.9 | 9.9 | 11.6 | 1.2 | 0.7 | 3.2 | 4.8 |
| JB Diego | 110.7 | 113.1 | 10.4 | 11.0 | 0.4 | 1.6 | 3.8 | 5.9 |
| KWS Santiago | 105.5 | 115.5 | 10.3 | 12.1 | 0.7 | 1.6 | 2.8 | 4.6 |
| Maris Wigeon | 119.0 | 113.7 | 10.1 | 10.1 | 1.0 | 4.6 | 3.7 | 5.1 |
| Milns N 59 | 101.9 | 83.1 | 9.1 | 11.2 | 0.8 | 2.1 | 3.6 | 5.1 |
| MV Kolo | 100.5 | 121.3 | 10.2 | 11.0 | 1.6 | 0.6 | 2.8 | 4.7 |
| Ostka Skomoroska | 107.0 | 93.9 | 8.8 | 9.8 | 1.4 | 1.7 | 3.7 | 4.9 |
| Red Lammas | 93.1 | 107.7 | 8.4 | 11.7 | 1.9 | 2.3 | 6.0 | 5.0 |
| Red Standard | 87.9 | 94.1 | 11.2 | 12.4 | 1.7 | 3.5 | 4.8 | 5.4 |
| Red Stettin 13 | 94.0 | 89.4 | 12.1 | 11.4 | 1.9 | 3.3 | 8.0 | 5.3 |
| Robigus | 99.3 | 125.8 | 10.8 | 10.4 | 0.7 | 2.1 | 3.4 | 5.1 |
| Samanta 117 | 98.6 | 121.4 | 8.2 | 8.3 | 0.9 | 0.3 | 2.5 | 3.8 |
| Soissons | 98.1 | 113.4 | 8.4 | 13.1 | 0.5 | 1.2 | 2.7 | 5.2 |
| Steadfast | 112.7 | 108.1 | 10.8 | 11.7 | 2.0 | 3.6 | 2.0 | 5.5 |
| Tiepolo | 118.2 | 111.3 | 11.2 | 12.0 | 1.1 | 2.0 | 2.2 | 4.3 |
| WW 502 | 117.2 | 95.2 | 8.8 | 8.7 | 0.9 | 1.0 | 2.0 | 5.3 |
| WW 512 | 104.8 | 89.7 | 14.2 | 10.4 | 1.0 | 2.6 | 3.6 | 4.4 |
| Mean | 105.2 | 108.0 | 10.0 | 11.0 | 1.1 | 2.0 | 3.5 | 5.0 |
| Standard errors of differences between means |  |  |  |  |  |  |  |  |
| Average | 10.24 | 9.65 | 1.09 | 1.09 | 1.36 | 1.38 | 1.26 | 0.53 |
| Maximum | 10.56 | 10.94 | 1.10 | 1.10 | 2.73 | 2.04 | 1.33 | 0.57 |
| Minimum | 10.20 | 9.42 | 1.07 | 1.08 | 1.22 | 1.22 | 1.19 | 0.53 |

Significant genotype, year and genotype-by-year interaction effects were found for SRN (Table 3), whereas no significant effects of tillage were found on SRN.

### 3.2. Effects of seeding rate and genotype interactions

Trials at the Duxford site investigated the effects of increased seeding rate on root phenotypes and genotypic responses to these effects. There were significantly fewer crown roots per plant at the higher rate (11.5) than standard rate (13.6). Yield was also greater at the higher seeding rate (8.5 t/ha) than standard rate (8.1 t/ha) (Table 4). However, there was no effect of sowing rate on RA or SRN. Whilst the main effect of sowing rate on NRN was non-significant, a highly significant genotype-by-sowing-rate interaction effect on NRN was found (Table 4). When the data for each seeding rate were analysed separately, highly significant differences were found among genotypes at both standard seeding rate (Wald statistic/d.f. = 3.69, P<0.001) and high seeding rate (Wald statistic/d.f. = 4.78, P<0.001). When the effect of sowing rate was analysed separately for each variety, varieties such as ‘Slejpner’ and ‘Flamingo’ had significantly fewer nodal roots at higher seeding rate (P<0.01 for both varieties) than standard rate, whereas ‘Robigus’ had significantly more nodal roots at the higher seeding rate (P<0.05).

## 4. Discussion

There has been increasing interest in investigating crop root phenotypes, especially in relation to resource use efficiencies and sustainability. We employed the field phenotyping method of shovelomics to characterise wheat root phenotypes in two sets of diverse wheat accessions, including landraces, historic and modern cultivars, to investigate changes in wheat root phenotypes due to breeding as well as the effects of crop management practices of tillage and sowing rate.

### 4.1. Temporal changes in wheat root traits

Correlating root traits against the year of variety release in the Duxford dataset revealed that whilst yields have linearly increased by approximately 0.04 t/ha/year, which is similar to 0.07 t/ha/year trends found by Mackay et al. (2011), this has been accompanied by a decline in numbers of crown and particularly nodal roots, as well as, to some extent, a widening of root angles. This trend in the Sonning dataset is particularly strong, where the varieties extended to pre 20^th^ century material, and suggests that the effect is due to continuous selection for yield over long periods rather than the rapid introduction of dwarfing genes in the 1960s. Other studies have found similar changes in root traits over time (Waines and Ehdaie, 2007). This finding reflects long-term trends in which crop plants have been selected to be less selfish and competitive as individuals (Denison 2012; Donald 1968). Early crop plants grown in heterogeneous stands may have had larger root systems due to natural selection for traits that allowed individual plants to usurp resources from their neighbours. However, continuous selection for crop genotoypes that are collectively more productive (a form of group-level selection) is expected to favour root traits that make individual plants less selfish (Zhu et al., 2019b). This is supported by recent work finding that higher crop yields of modern wheat varieties are associated with reduced root numbers (Zhu et al., 2019a).

Our study also found that RA increased over time in the set of varieties tested at Sonning. RA has been identified as an important adaptive trait to water-limited environments, where genotypes with a narrower angle are able to access water at greater depths (Manschadi et al., 2006). Lynch et al. (2007) also suggested a strategy of selection of ‘steep, cheap and deep’ roots for improved adaptation of maize to water limited environments. Our results suggest that whilst a narrower root angle may be beneficial for crop adaptation in water limited environments, this has not been the direction of breeders’ selection in UK winter wheat where modern elite varieties exhibit a wider angle than do older UK varieties. This may be because of the complexity of environmental and agronomic factors affecting yield in the UK (Mackay et al., 2011), and so possibly may have more to do with agronomy than just water availability, which is likely the case in drier areas.

Intensification of agriculture and increased fertiliser use (Glass, 2003) could also explain the reduction in NRN in modern UK varieties. It has been suggested that lower root densities in the upper soil profile, but which extend to a greater depth, are required for efficient uptake of nitrate, which is made readily available and mobile in soil due to synthetic fertiliser application (Lynch, 2013; White et al., 2013). On the other hand, the value of an RSA characterised by increased root number and at a shallower angle has been found to be particularly important for scavenging and uptake of phosphorus, which is relatively immobile in soil and more abundant and available in the upper soil profile (Lynch and Brown, 2001; Péret et al., 2014). Therefore, a trade-off potentially exists for uptake of these two key nutrients, which differ in spatial and temporal distribution and availability within the soil profile according to production system and soil management regime. For example, in non-inversion tillage systems, soil organic matter and associated phosphorus is often stratified and concentrated in the topsoil (Poirier, 2009). Manske and Vlek (2002) advocate a high-input root ideotype characterised by seminal root dominance in contrast to a low-input ideotype based on a greater number of roots to explore the soil volume. Increasing root number is also thought to increase crop plant competitive ability against weeds (Richards, 2007) which are particularly problematic in low-input environments (Hoad et al., 2012). Our results support this, demonstrating that modern elite varieties, which are adapted to high-input environments, have a smaller number of nodal roots. We suggest that utilisation of historic cultivars as breeding material would be useful to improve the adaptation of modern varieties adapted to environments where nutrients are not made readily available through application of inorganic fertilisers, or where reductions in input use is a priority.

### 4.2. Effects of tillage

Significant effects of tillage on three of the measured root traits (RA, CRN, NRN) suggests a general sensitivity of RSA to the growing environment. Numerically, the difference appears small, but small differences in RA can result in a larger spread of the root system at depth. In addition, significant genotype by tillage interactions for these traits suggests that this sensitivity is genotype specific. Consistent genotype effects on RA across treatments were only found in year 1 in the CT system. These interactions underline the importance of understanding and reporting soil management practices for fields used in root phenotyping experiments. Inversion tillage in the CT system, which would likely cause smaller soil bulk density in the upper profile than non-inversion tillage (Tebrügge and Düring, 1999), likely provides a better environment for maximising and observing genotypic differences in RA. Genotype by tillage interactions for CRN and NRN in both years indicate that the production of crown and nodal roots by different genotypes also depends on soil management. There were fewer nodal roots produced in SNI than CT, and grain yield was also lower in SNI than CT in both years (personal communication). These results corroborate findings that reduced yields are often found in SNI practices (Pittelkow et al., 2015). These genotype-by-tillage interactions also suggests that selection of genotypes in the target environment would be required in order to improve adaptation to conservation agriculture systems characterised by reduced or non-inversion tillage. This would enable enhanced performance with reduced tillage, as part of conservation agriculture systems, which are able to make more efficient use of nutrients (Habbib et al., 2016).

### 4.3. Effects of seeding rate

Trials at the Duxford site compared a diverse set of wheat varieties at standard and high seeding rates. This enabled the investigation of varying genotypic responses in root traits to increased density and within crop competition. Our results found that higher densities generally decreased CRN, but that the effect on NRN was highly genotype specific with some varieties responding positively but some negatively. This effect of reduced CRN closely reflects results in barley reported by Hecht et al. (2019) where root numbers, together with tiller number, declined at higher densities. However, this is contrary to results of O’Brien et al. (2005) who found an increase in pea root proliferation with increased competition but with equal nutrient availability per plant. Hecht et al. (2016) also found an increase in root density from fine root branching as a response to increased density, which suggests independent control of crown root numbers and root branching. Our results showing reduced CRN suggests that this is a result of limited nutrient availability due to increased competition at higher densities rather than an adaptive response to competitors. The competitive and compensatory relationships among crop plants and tillers on the same plant are well known (e.g. Nerson, 1980). As yields were found to be significantly higher at sowing rates well above the standard practice in the study presented here, adaptation of crop varieties to higher densities would be an opportunity for yield improvement. However, significant genotype-by-sowing-rate interactions were only found for NRN and not yield in the Duxford dataset. Therefore, there is no evidence here that varieties which respond differently to density in terms of NRN are able to yield more at higher densities. It may be hypothesised that the more modern varieties would exhibit a less competitive response to increased density and produce fewer nodal roots, as outlined above in relation to selection for decreased intra-crop competitive effects (Zhu et al., 2019b). However, we found no relationship between NRN response to selection and variety release date, and therefore, the implications of this genotype-by-sowing rate interaction remain unclear. No effect of seeding rate on RA or SRN was found which may be because of the greater variability of these traits. However, more vertical root angles in response to competition were found in a study in maize (Shao et al., 2018), which suggests biological effects exist but were not detected in the present study.

### 4.4. Application of shovelomics

Our ability to detect genotypic differences in RSA confirms that shovelomics is an effective method to phenotype wheat root traits in the field, corroborating a recent study in wheat (York et al., 2018). However, here we also investigated effects of management practices including contrasting tillage system and increased sowing rate. Whilst classification of cereal root types are rarely standardised (Zobel and Waisel, 2010) and crown and nodal roots are often considered together (Manske and Vlek, 2002; York et al., 2018), we were able to differentiate between these root classes finding clear genotypic differences, particularly in NRN. Although the method only observes roots present in upper soil layers, the advantage is that roots are sampled *in situ*, in a real field environment, unlike pot- or pipe-based root phenotyping systems in which expression of root traits are likely affected by the container and the nature of the rooting medium (Passioura, 2006). Time requirements are an important consideration in root phenotyping. We found sample collection to take approximately 2 minutes per experimental plot with subsequent washing taking between 5 to 10 minutes and imaging taking approximately 0.5 to 2 minutes per sample. Up to 20 wheat genotypes were characterised in each environment under multiple treatments in this study, but greater throughput would be required for marker discovery using genetic mapping populations or screening lines in a wheat breeding programme. However, the shovelomics method could be used to identify desirable root phenotypes in novel germplasm that could be integrated into pre-breeding programmes, or to validate genetic effects found in controlled environment phenotyping methods. The method does not provide information on root traits in deeper soil layers; for this, soil coring (e.g. Wasson et al., 2016) or other methods are required.

### 4.5. Conclusions

In summary, we found significant genotypic variation for RSA phenotypes, the expression of which differed according to the tillage regime, sowing rate and growing environment. Our results suggest that selective breeding for yield has resulted in a reduction in later developing root numbers, in particular nodal roots. The results raise new questions about the role of tillage regime and sowing density on root traits, but further research is required to understand which combination of root traits are most beneficial for a given environment or soil management scenario. The information about differences in RSA traits identified here can contribute to improving crop adaptation by matching specific root traits to specific target environments or crop and soil management practices. In future work, questions should be addressed such as how tillering capacity and CRN are related and interact with stand density, and the nature of trade-offs between RA, lodging susceptibility, and growth under varying levels of nitrogen inputs.

## Glossary

CRN: Crown root number
CT: Conventional tillage
NRN: Nodal root number
RA: Root angle
RSA: Root system architecture
SNI: Shallow non-inversion tillage
SRN: Seminal root number

## 5. Acknowledgements

The authors acknowledge funding for the trials at the Sonning site from the European Community’s Seventh Framework Programme (FP7/ 2007-2013) under the grant agreement no. FP7-613556 (WHEALBI); the Nuffield Foundation for funding work of GE and the Biotechnology and Biological Sciences Research Council (BBSRC) under grant agreement BB/M011666/1 and BB/M013995/1 for funding trials at the Duxford site. We thank farm staff at the University of Reading, Sonning farm field trial site; Ambrogio Costanzo and Dominic Amos of The Organic Research Centre; the Trials Team at NIAB Cambridge for management of field trials; Simon Orford of the John Innes Centre Germplasm Resource Unit for providing seed. Greg Mellers and Ian Mackay gave invaluable advice on statistical analysis.

## 6. Author contributions

EO, HJ, JC, JB and NF conceived the work and provided supervision, GE, HJ, EM, AO and NF performed sample collection and analysis, NF analysed the data, NF and EM wrote the paper and all authors edited the manuscript.

## 8. Supplementary Material

**Supplementary table 1.**
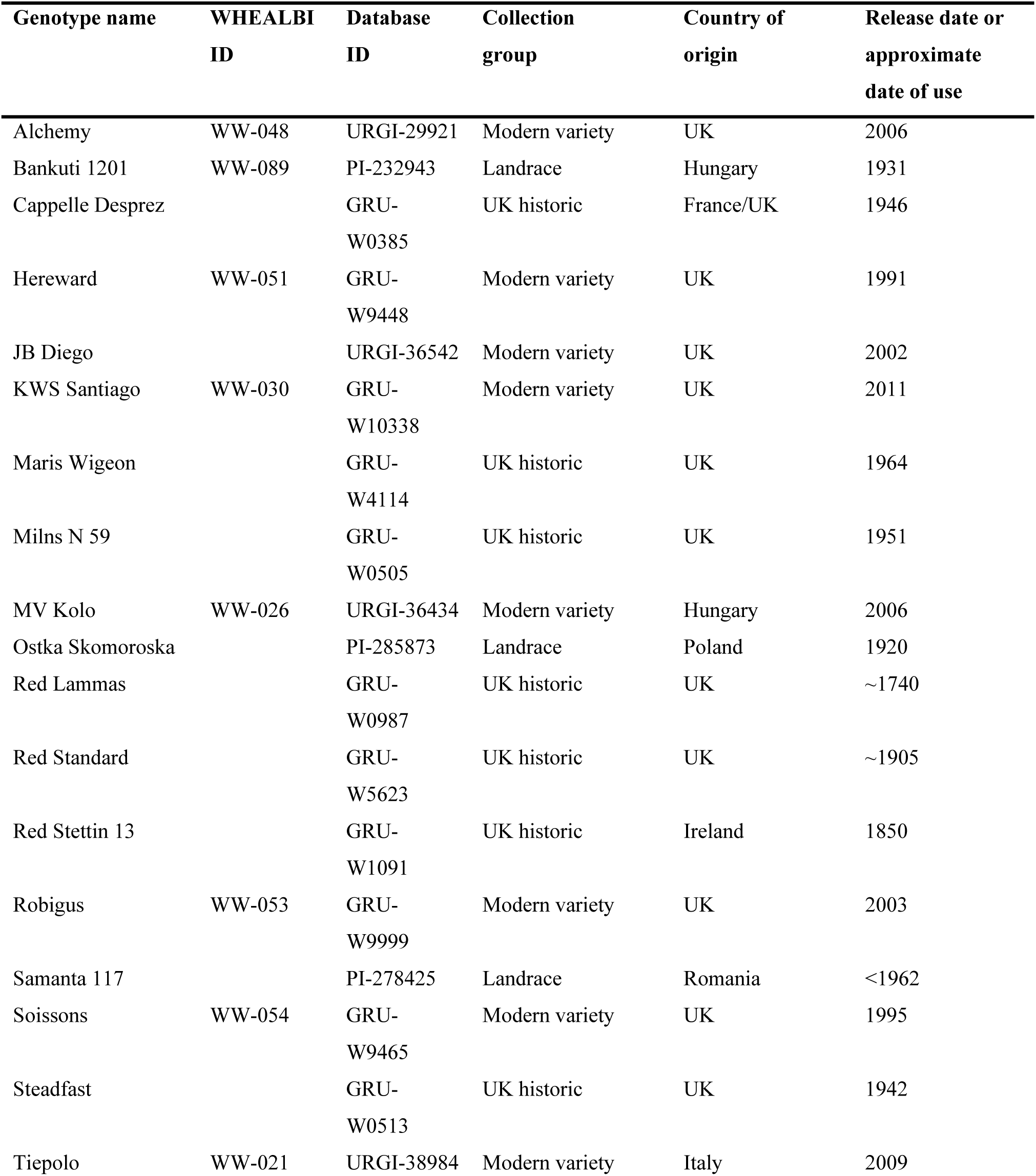
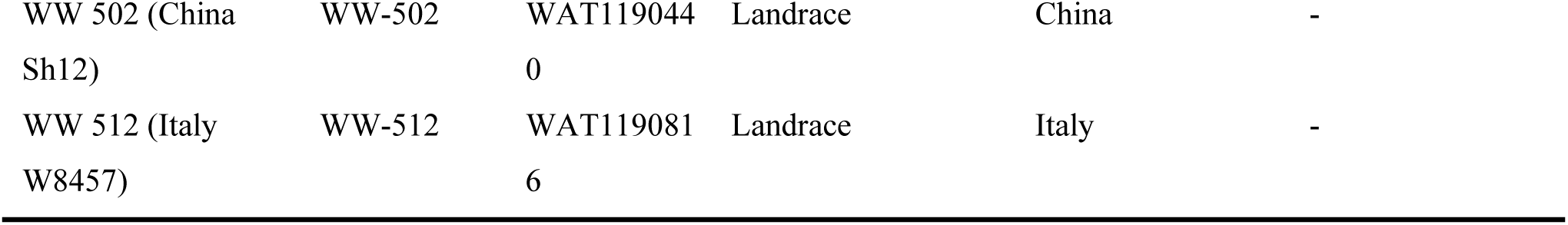
Origins and dates of winter wheat cultivars as part of the WHEALBI panel. Dates of use for some genotypes are unavailable. Database ID with prefixes ‘PI’ are from the USDA GRIN database (https://npgsweb.ars-grin.gov/); ‘GRU’ and ‘WAT’ from the Germplasm Resource Unit and Watkins Collection (https://www.seedstor.ac.uk); ‘WW’- are WHEALBI accessions (http://www.whealbi.eu/); ‘URGI’ from GnpIS (https://urgi.versailles.inra.fr/gnpis/).

**Supplementary table 2.** Origins and release dates of the 16 wheat varieties that represent the founders of the NIAB Diverse MAGIC population.

| Name | Country of origin | Adaptation | Release date |
| --- | --- | --- | --- |
| Banco | Sweden | Winter | 1956 |
| Bersee | UK/France | Winter | 1951 |
| Brigadier | UK | Winter | 1993 |
| Copain | France | Winter | 1980 |
| Cordiale | UK | Winter | 2004 |
| Flamingo | NL/DK | Winter | 1960 |
| Gladiator | UK | Winter | 2004 |
| Holdfast | UK | Winter | 1935 |
| Kloka | Germany | Facultative | 1965 |
| Maris Fundin | UK | Winter | 1975 |
| Robigus | UK | Winter | 2003 |
| Slejpner | Denmark/Sweden | Winter | 1986 |
| Soissons | France | Winter | 1995 |
| Spark | UK | Winter | 1993 |
| Steadfast | UK | Winter | 1942 |
| Stetson | UK | Winter | 1983 |

**Supplementary table 3.** Soil chemistry measurements at the Sonning site in both years.

|  | Year 1 | Year 2 |
| --- | --- | --- |
| pH | 6.2 | 6.0 |
| NO<sub>3</sub> -N (mg kg<sup>-1</sup>) | 4.08 | 0.98 |
| P (mg l<sup>-1</sup>) | 30.6 | 40.4 |
| P Index | 3 | 3 |
| K (mg l<sup>-1</sup>) | 59.6 | 196.0 |
| K Index | 0 | 2+ |
| Mg (mg l<sup>-1</sup>) | 37.6 | 68.0 |
| Mg Index | 1 | 2 |
| Organic matter (%) | 2.6 | 2.4 |

**Supplementary table 4.** Raw data per plot used for analysis from both the Duxford and Sonning trials sites.

